# Evolution of Mating Types Driven by Purifying Selection against Mitochondrial Mutations

**DOI:** 10.1101/049577

**Authors:** Arunas L Radzvilavicius

**Affiliations:** Department of Genetics, Evolution and Environment & CoMPLEX; University College London, Gower Street, WC1E 6BT London, United Kingdom

**Keywords:** mating types, mitochondria, uniparental inheritance, purifying selection, sexes

## Abstract

Sexual cell fusion combines genetic material of two gametes, but why the two reproductive cells have to belong to distinct self-incompatible gamete classes is not known. In a vast majority of sexual eukaryotes, mitochondria are inherited uniparentally from only one of the two mating types, which is thought to facilitate purifying selection against deleterious mitochondrial mutations and limit the inter-genomic conflicts. Here I argue that two mating types in eukaryotes represent a mechanism of mitochondrial quality control through the highly asymmetric transmission of mitochondrial genes at cell fusion. I develop a mathematical model to explicitly study the evolution of two self-incompatible mating type alleles linked to the nuclear locus controlling the pattern of organelle inheritance. The invasion of mating-type alleles is opposed by the short-term fitness benefit of mitochondrial mixing under negative epistasis and the lower chance of encountering a compatible mating partner. Nevertheless, under high mitochondrial mutation rates purifying selection against defective mitochondria can drive two mating types with uniparental inheritance to fixation. The invasion is further facilitated by the paternal leakage of mitochondria under paternal control of cytoplasmic inheritance. In contrast to previous studies, the model does not rely on the presence of selfish cytoplasmic elements, providing a more universal solution to the longstanding evolutionary puzzle of two sexes.

## Introduction

Sex is among the traits universal to all complex life and was therefore already present in the last eukaryotic common ancestor. By combining the genetic material of two gametes, sexual cell fusion and meiotic recombination exposes the hidden genetic variation in finite populations, breaks up the unfavorable allelic combinations under fluctuating selection, and mitigates the mutational meltdown (Otto, 2009). In a vast majority of cases, the two mating partners belong to distinct self-incompatible gamete classes, i.e. mating types in isogamous protists or true sexes with anisogametes, but the selective forces behind this fundamental asymmetry remain elusive (Billiard et al., 2010). The existence of two gamete types in most eukaryotes is often regarded as an evolutionary conundrum, as it reduces the number of potential mating partners, which should be detrimental if the cost of finding a compatible gamete is high and the mating opportunities are limited.

Several non-exclusive explanations for the emergence of self-incompatible gamete types in a unisexual population have been proposed over the years (Billiard et al., 2010). Two mating types might have appeared together with bipolar gamete-recognition systems ensuring efficient intercellular signaling (Hoekstra 1982; Hadjivasiliou et al., 2015), to promote outbreeding (Charlesworth and Charlesworth, 1979, Uyenoyama, 1988) or to improve the mitochondrial-nuclear coadaptation (Hadjivasiliou et al., 2012). Particularly appealing has been the idea that mating types emerged to ensure the asymmetric inheritance of cytoplasmic genetic elements. Indeed, in a vast majority of eukaryotes, only one gamete class transmits its organelles— mitochondria and chloroplasts—to the zygote, although the uniparental inheritance is not always complete, with paternal leakage and heteroplasmy being relatively common (Breton and Stewart, 2015). Early theoretical studies supported the hypothesis, but relied on the presence of the so-called “selfish” or parasitic cytoplasmic elements, simplistic assumptions, and lacked generality or empirical support (Hastings, 1992; Hurst and Hamilton, 1992; Hutson and Law, 1993).

A more general view of the evolutionary advantage of uniparental inheritance (UPI) is that it improves the efficacy of purifying selection against mitochondrial mutations (Bergstrom and Pritchard, 1998; Hadjivasiliou, 2013), and therefore confers a long-term fitness advantage. Mitochondrial mixing, on the other hand, reduces the strength of selection and is costly in a long term (Radzvilavicius, 2016). Asymmetric transmission of mitochondria therefore counters the mutational meltdown in mitochondrial genomes, at least partially accounting for the highly efficient purifying selection in the absence of recombination (Cooper et al., 2015). A recent study explicitly accounting for the segregation of detrimental mitochondrial mutations, however, found that the fitness benefit of UPI decreases in a frequency-dependent manner, and concluded that UPI alone is unlikely to drive the evolution of two self-incompatible mating types in an ancestral unisexual population (Hadjivasiliou, 2013). The robustness and generality of these results, however, is limited by the assumptions of the study; in particular, the peculiar mating kinetics in which gametes unable to find a suitable mating partner immediately incur a severe fitness cost are not representative of biological reality, and put the self-incompatible gametes at a strong disadvantage. Whether the asymmetric inheritance of mitochondria can facilitate the evolution of binary mating types under the mitochondrial mutation pressure remains unclear.

Here I develop a mathematical model to investigate whether selection against deleterious mitochondrial mutations can drive the evolution of two self-incompatible mating types by establishing the asymmetric transmission of mitochondrial genes. Mitochondria continuously accumulate mutations which are purged by selection on the level of the cell, with random mitochondrial drift generating additional variance at every cell division. Nuclear inheritance-restriction alleles regulate the selective destruction of mitochondria inherited either from the cell carrying the allele, or its mating partner in the opposite nuclear state. Mating kinetics play a central role in the model, with long mating periods and low gamete mortality rates favoring the invasion of self-incompatible gamete classes. Strikingly, paternal leakage of mitochondria—an evolutionarily stable state under paternal control of cytoplasmic inheritance—can facilitate the evolution of binary mating types under negative epistasis between mitochondrial mutations.

## Modelling the evolution of uniparental inheritance

### Two modes of UPI regulation

Mechanisms of uniparental inheritance vary substantially across eukaryotic species (Sato and Sato, 2013), but one general theme seems to be rather common: one of the mating partners tags its mitochondria with the mating-type specific marker protein, which is recognized by the partner’s molecular factors after the gamete union, leading to the eventual degradation of the marked organelles. In mammals, for example, sperm mitochondria are tagged by the recycling marker protein ubiquitin and destroyed by the egg’s cytoplasmic destruction machinery after the gamete union (Sutovsky et al., 1999). Similarly, in basidiomycete yeast *Cryptococcus neoformans*, genes *SXI1a* and *SXI2α* located in opposite mating types are responsible for tagging and recognition of paternal mitochondria (Yan et al., 2007).

In the present work I assume that uniparental inheritance can be established in two ways, based on how the two gametes control organelle transmission (Fig. 1). First, a cell (wild-type allele *B)* can develop the ability to recognize a universal mitochondrial marker protein, at the same time protecting its own organelles from degradation, e.g. by ceasing the expression of the marker in its own mitochondria (allele *U*_*m*_, Fig. 1b). I term this mechanism the “maternal” mode of UPI, and the gamete destroying its partner’s mtDNA the “maternal” gamete—the definition which does not rely on the prior existence of mating types or sexes and merely reflects the fact that the gamete controlling the cytoplasmic inheritance eliminates its partner’s mtDNA, but not its own (Fig. 1b). Alternatively, a cell can start producing a new nuclear-coded and universally recognized mitochondrial marker, but lose the ability to recognize the tag in its own cytoplasm (allele *U*_*p*_). In this case the gamete essentially controls the inheritance of its own organelles, as stopping the expression of the mitochondrial tag protein would make them unrecognizable in the zygote. I term this mechanism the “paternal” mode of UPI (Fig. 1c). In gamete unions with identical alleles at the mitochondrial-inheritance locus, both mating partners lack either the mitochondrial marker or the corresponding molecular destruction machinery. The inheritance of mitochondria is therefore biparental in *U*_*m*_×*U*_*m*_ and *U*_*p*_×*U*_*p*_ gamete unions.

**Figure 1.**
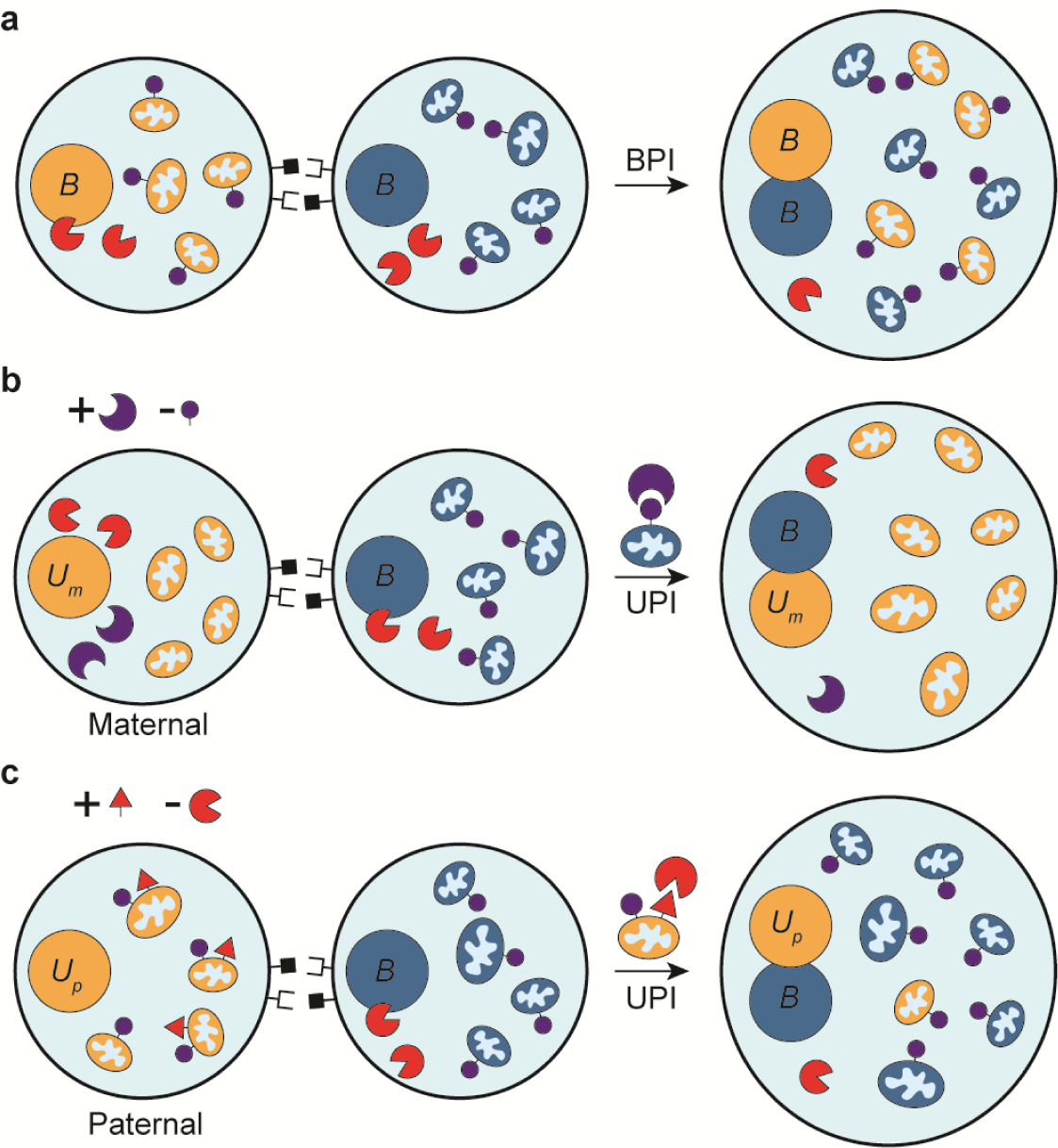
Two modes of uniparental inheritance differ in the way organelle inheritance is controlled. In an ancestral state, mitochondrial surface proteins do not have corresponding nuclear-coded molecular factors targeting the organelles for destruction (**a**). Maternal mode of UPI arises with one of the gametes developing the ability to recognize the mitochondrial marker protein universal to the whole population, while protecting its own organelles e.g. by removing the tag (**b**). Similarly, in paternal mode of UPI, one of the gametes marks its mitochondria with a new universally recognizable marker, but loses the ability to target it for destruction in its own cytoplasm (**c**). Alternatively, the paternal gamete can simply remove a part of its own organelle population before fertilization.

### Mating kinetics

The rules governing the frequency of gamete unions of distinct mating types are expected to play a critical role in the origin and evolutionary stability of mating types (Iwasa and Sasaki, 1987). Rare sexes, for instance, are favored in models where mating opportunities are limited and only a short period of time is available to locate a suitable mating partner, but the same models penalize the newly emerging self-incompatible gamete classes in unisexual populations where all gametes are initially compatible. A more realistic scenario should assume that the mating period can be significantly longer than the duration of a single cell-fusion attempt, and so the gametes can wait for a suitable mating partner to arrive before being eliminated from the population.

In this work I adopt the mating kinetics first developed in the models of Iwasa and Sasaki (1987), and Hutson and Law (1993). I assume the gamete pool of finite size, in which the influx of gametes matches their removal due to random death and zygote formation. The gamete influx rates are proportional to the allele frequencies within the infinite population, while the gamete death rate is kept at a constant level *δ*. Cells within the gamete pool form random pairwise associations at rate *β/2*, and are removed from the pool if they are compatible and able to fuse. The allele frequencies in the next generation are calculated from the steady-state gamete-class frequencies within the mating pool. These frequencies correspond to the equilibria of the following system of equations

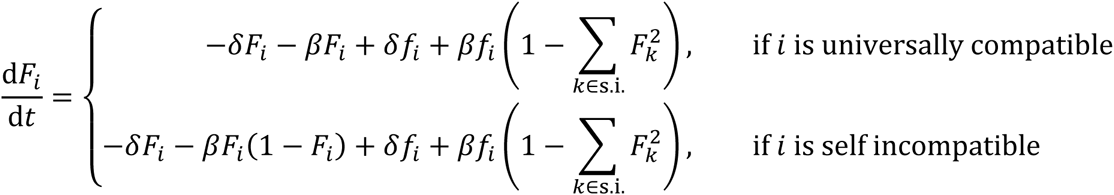

Here *F*_*i*_ denotes the frequency of the genotype *i* in the gamete pool, while *ƒ*_*i*_ is the corresponding frequency within the infinite population. The sums here are over the self-incompatible gamete classes (s.i.). I fix the mating rate at *β* = 1 and vary only the value of *δ*, as the steady-state frequencies of gamete classes depend only on the ratio of these rates *β/δ*.

### Population life cycle

The model assumes an infinite population of unicellular haploid organisms, each containing *M* mitochondria, and is in many ways similar to the model developed in (Radzvilavicius, 2016). The population state can be represented by the (*M* + 1) × *n* matrix **P**, where the matrix element *P*_*i,j*_ denotes the frequency of cells in a nuclear state *j* ≤ *n* and containing *m* mutant mitochondria. The horizontal index *j* enumerates all possible nuclear states including the mating type and the mode of mitochondrial transmission.

Mitochondrial mutation is represented by the transition **P**^(*t*,1)^ = **UP**^(*t*)^, where the matrix element *U*_*i,j*_ represents the probability that a cell with *j* mutant mitochondria will have *i* mutants after the mutation event,

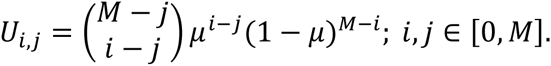

Mutation occurs in individual mitochondria, but selection acts on the level of the cell, i.e. on groups of mitochondria. In our model, cell fitness *w* directly depends only on the mitochondrial part of cell’s genotype. The updated population state after selection is then

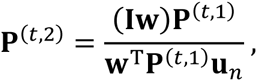

where **I** is the identity matrix. **u**_*n*_ is a column vector of ones, so that **P**^(*t*, 1)^**u**_*n*_ is the row-wise sum of **P**^(*t*, 1)^ and **w** is a column vector containing all possible values of mitochondrial fitness,

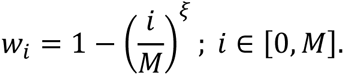

Parameter *ξ* determines the magnitude of epistatic interactions between mitochondrial mutations. With ξ > 1 the fitness cost of every new mutation increases with the overall mutation load (negative epistasis). Negative epistasis between deleterious mitochondrial mutations leads to mitochondrial threshold effects, well known from empirical observations in eukaryotes (Rossignol et al., 2003).

Gametes in the gamete pool fuse at random, according to their equilibrium frequencies *F*_*i*_. For asymmetric gamete unions, assuming that the gamete *k* is of maternal type, we have,

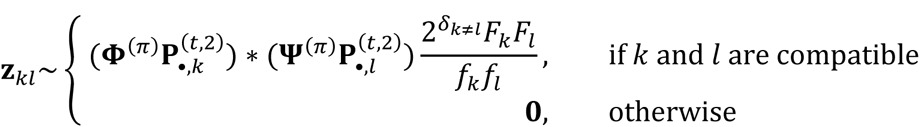

An asterisk here denotes vector convolution, *ƒ*_*k*_ is the frequency of cells in a nuclear state *k*, i.e. 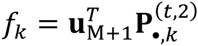 and *F*_*k*_ is the corresponding equilibrium frequency in the gamete pool. The delta symbol *δ*_*k ≠ l*_ = 1 if *k ≠ l* and is 0 otherwise. The zygote-state vectors **z**_*kl*_ are scaled linearly to sum up to one. The two transition matrices **Φ**^(π)^ and **Ψ**^(π)^ are included to implement the mitochondrial inheritance bias where one of the gametes transmits more mitochondria than the other. We assume that the paternal gamete contributes *πM* mitochondria through sampling without replacement, and that (2 - *π)M* mitochondria come from the maternal gamete via random sampling with replacement 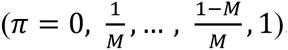. The two transition matrices therefore have elements

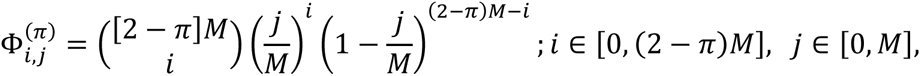

and

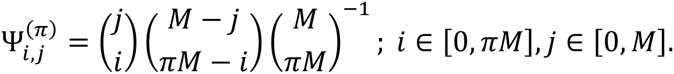

The life cycle is completed by two meiotic cell divisions restoring the haploid state. At the start of the next generation the genotype frequencies then are

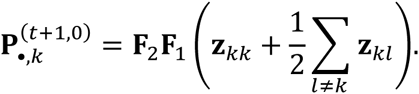

I do not differentiate between **z**_*kl*_ and **z**_*lk*_, i.e. both state vectors indicate the same zygote type. **F**_1_ and **F**_2_ are transition matrices for the two meiotic divisions implemented as mitochondrial sampling without replacement. Their corresponding elements are

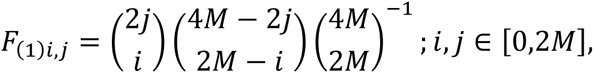

and

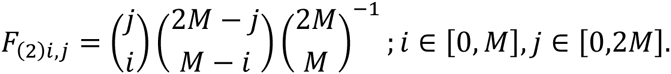

The following results are based on the numerical solution of the above system of equations.

## Asymmetric sex before mating types: UPI facilitates purifying selection against deleterious mitochondrial mutations

Eukaryotic sex depends on molecular mechanisms for self/non-self-recognition and gamete attraction (Goodenough and Heitman, 2014); similar mechanisms likely existed in both the bacterial ancestor of mitochondria and the archaeal host in the form of quorum-sensing and biofilm-formation systems triggered by external conditions (Ng and Bassler, 2009; Fröls, 2013). With the prokaryotic ancestry of eukaryotic gamete attraction machineries, the initial proto-eukaryotic self-recognition system was likely symmetrical, i.e. initial forms of sexual reproduction did not involve differentiation into distinct gamete classes (Goodenough and Heitman, 2014; Heitman, 2015). Self-incompatible mating types and sexes then could have been a later addition, driven, as I shall argue here, by purifying selection against mitochondrial mutations.

Assuming that the initial form of sexual reproduction was unisexual, I first model the evolution of uniparental inheritance in an ancestral population without mating types. While all pairs of gametes are capable of mating, only the unions between *U*_*m*_ and *B* (or *U*_*p*_ and *B)* involve the asymmetric transmission of mitochondria (Fig. 1). As I show in Fig. 2, both nuclear alleles *U*_*m*_ and *U*_*p*_ invade and can reach the frequency of *e*_2_ = 0.5, but the invasion dynamics differ substantially between the two modes of UPI control.

**Figure 2.**
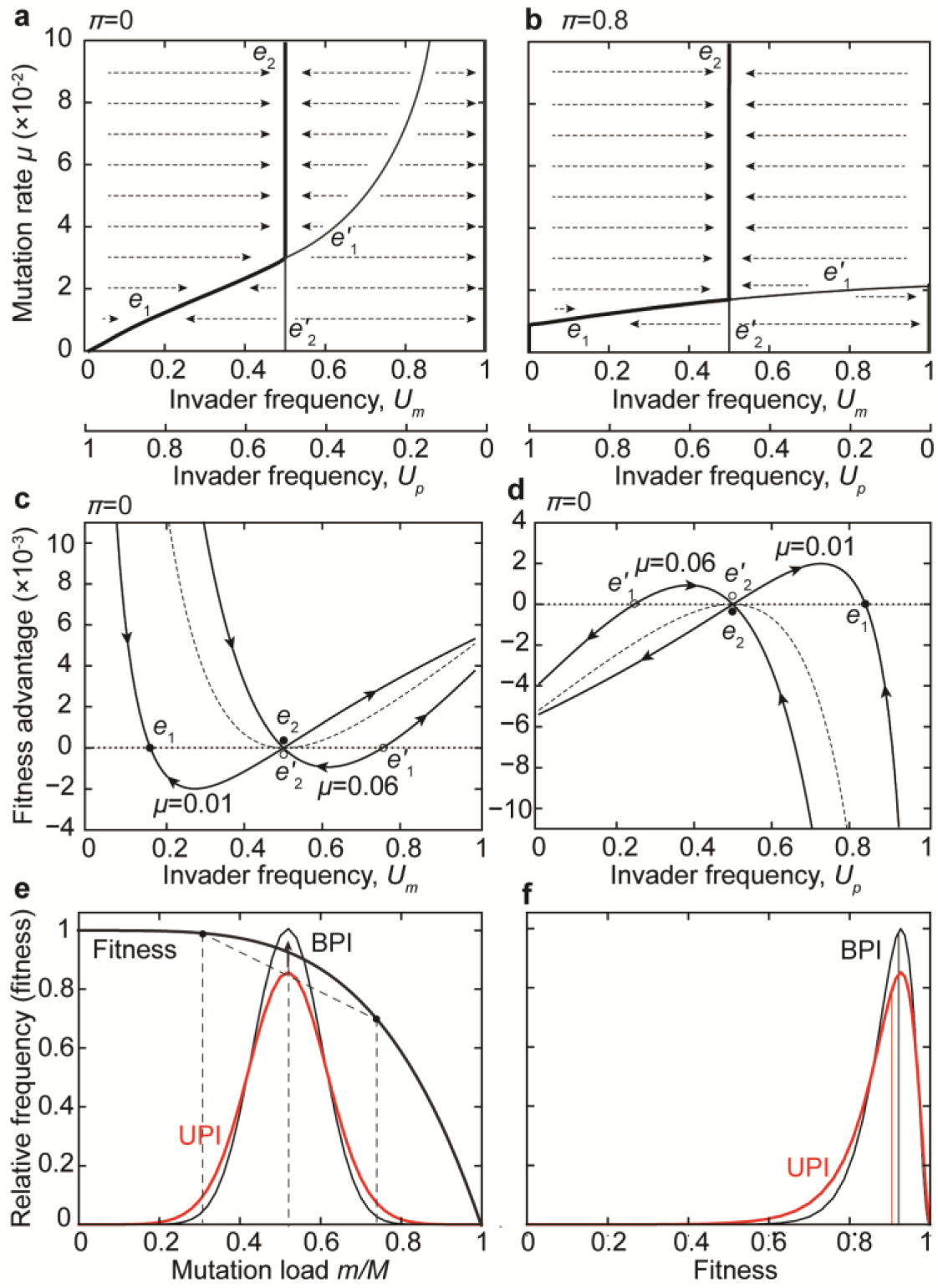
Invasion of the uniparental inheritance alleles in a population without mating types. Equilibria for the maternal *(U*_*m*_) and paternal *(U*_*p*_) UPI alleles for strict uniparental inheritance *π*=0 (**a**) and asymmetric inheritance with mitochondrial leakage *π*=0.8 (**b**). Equilibria *e*_1_ and *e*_2_ (thick lines) are stable, while *e'*_1_ and *e'*_2_ indicate unstable states. Fitness advantage of the UPI mutants *U*_*m*_ (**c**) and *U*_*p*_ (**d**) with *π* =0 depends on their frequency, explaining the stability of equilibria in (a). Dashed line indicates the fitness landscape at the saddle point *μ* = 0.03, where equilibria *e*_1_ and *e*_2_ coincide. The strength of epistasis *ξ* = 2, *M* = 50. The asymmetric inheritance increases variance in the mutation load allowing for more efficient purifying selection and giving a long-term advantage, while mitochondrial mixing under the biparental inheritance (BPI) increases the frequency of intermediate cytoplasmic states (**e**). But with negative epistasis between mitochondrial mutations (**e**), mitochondrial mixing results in higher fitness than would be expected under linear fitness interactions (**f**), giving BPI a short-term advantage.

Starting at low allele frequencies, the invader *U*_*m*_ attains one of two distinct equilibria, *e*_1_ < 0.5 under low mutation rates and strong negative epistasis, or *e*_2_ = 0.5 under higher mutation rates (Fig. 2a). The analogous set of equilibria exists for the paternal invader *U*_*p*_, but this time *e*_2_ = 0.5 is the only asymmetric equilibrium which can be reached starting from low initial mutant frequencies (Fig. 2a). In this case the combination of the mutation rate and the initial frequency must exceed the unstable equilibrium 0 < *e'*_1_ < 0.5, the characteristic frequency of which approaches zero at high mutation rates. Paternal leakage of mitochondria relaxes the conditions for the invasion of *U*_*p*_ to the equilibrium *e*_2_ (Fig. 2b), but at the same time hinders the invasion of *U*_*m*_ under low mutation rates.

What determines the equilibrium frequencies of UPI alleles in a population without mating types? The model shows that the fitness advantage of *U*_*m*_ decreases as the allele invades from small frequencies (Fig. 2c), while the opposite is true for *U*_*p*_ (Fig. 2d), determining the locations of equilibria *e*_1_ and *e*_2_. This behavior can be explained in terms of costs and benefits of asymmetric transmission of mitochondrial genes, and the statistical association of these benefits to the nuclear allele of mitochondrial inheritance control.

Uniparental inheritance of mitochondria increases the frequency of extreme cytotypes underrepresented at the mutation-selection equilibrium, reducing the abundance of the common intermediate cytoplasmic states (Fig. 2e). This increase in mitochondrial variance facilitates purifying selection against deleterious mutations and gives the UPI mutant a long-term advantage over the biparental population. Nevertheless, under negative epistasis (Fig. 2e) intermediate cytoplasmic states have higher fitness than expected from the linear combination of the extremes, penalizing the UPI invader and giving the biparental inheritance a short-term benefit (Fig. 2f). These short-term fitness effects, however, are relevant only if the association between the cytoplasmic state and the nuclear allele of inheritance control is weak (e.g. with paternal leakage, when part of the mitochondrial population is inherited from the unrelated gamete); otherwise the long-term variance-based effects dominate. The stable equilibria *e*_1_ and *e*_2_ are located where short-term effects match the long-term fitness advantage of the UPI invader.

Since the rate at which the resident *B* inherits the cytoplasm from the uniparental mutant rises with increasing frequency of *U*_*m*_, the short-term fitness gains of *B* increase, halting the invasion at *e*_1_ < 0.5 or *e*_2_ = 0.5. Paternal leakage associated with *U*_*m*_ weakens the mito-nuclear associations and therefore reduces the strength of the long-term fitness effects. The opposite pattern of nuclear-cytoplasmic linkage applies to the invader with paternally-determined UPI: destroying part of one’s own mitochondria increases the level of transmission asymmetry, but weakens the statistical associations between the mitochondrial population and the allele *U*_*p*_. As the allele *U*_*p*_ invades from small frequencies, the rate of biparental unions *U*_*p*_ × *U*_*p*_ increases, but so does the overall frequency of uniparental transmissions. The short-term fitness gains of invading *U*_*p*_ therefore increase with its overall frequency. As the invasion proceeds, the biparental unions between the identical gametes become more common, eventually reversing the trend of frequency-dependence (Fig. 2c-d).

## Evolving paternal leakage in UPI

Suppose now that the level of paternal leakage *π* is itself an evolvable trait, controlled by a single nuclear allele *U*_*p*_ or *U*_*m*_. The amount of organelles transmitted to the progeny from each gamete can indeed be regulated genetically, e.g. by controlling the expression of gamete-specific protein markers tagging the organelles for selective degradation (e.g. ubiquitin). What pattern of organelle inheritance would we expect to evolve in the population without mating types? I performed evolutionary invasion analysis to find the evolutionarily stable states (ESS; Eshel, 1983; Geritz et al., 1998) for the paternal leakage *π* under both maternal and paternal modes of inheritance control.

With maternal regulation of cytoplasmic inheritance, the sole non-invadable state with asymmetric transmission of mitochondria is the strict UPI, i.e. *π*_ESS_ = 0 (Fig. 3a), at which the gamete *U*_*m*_ discards all mitochondria inherited from *B*. As the leakage *π* goes down, both the long-term fitness advantage of variance, and the strength of nuclear-cytoplasmic associations become more significant. The equilibrium frequency of *U*_*m*_ at *π*_ESS_ therefore increases with increasing mutant load, consistent with the role of UPI increasing the efficacy of purifying selection against mitochondrial mutations. Similarly, weak epistatic interactions (ξ → 1) reduce the short-term fitness gains in symmetric gamete unions, and result in higher equilibrium frequencies of *U*_*m*_ (Fig 3b).

**Figure 3.**
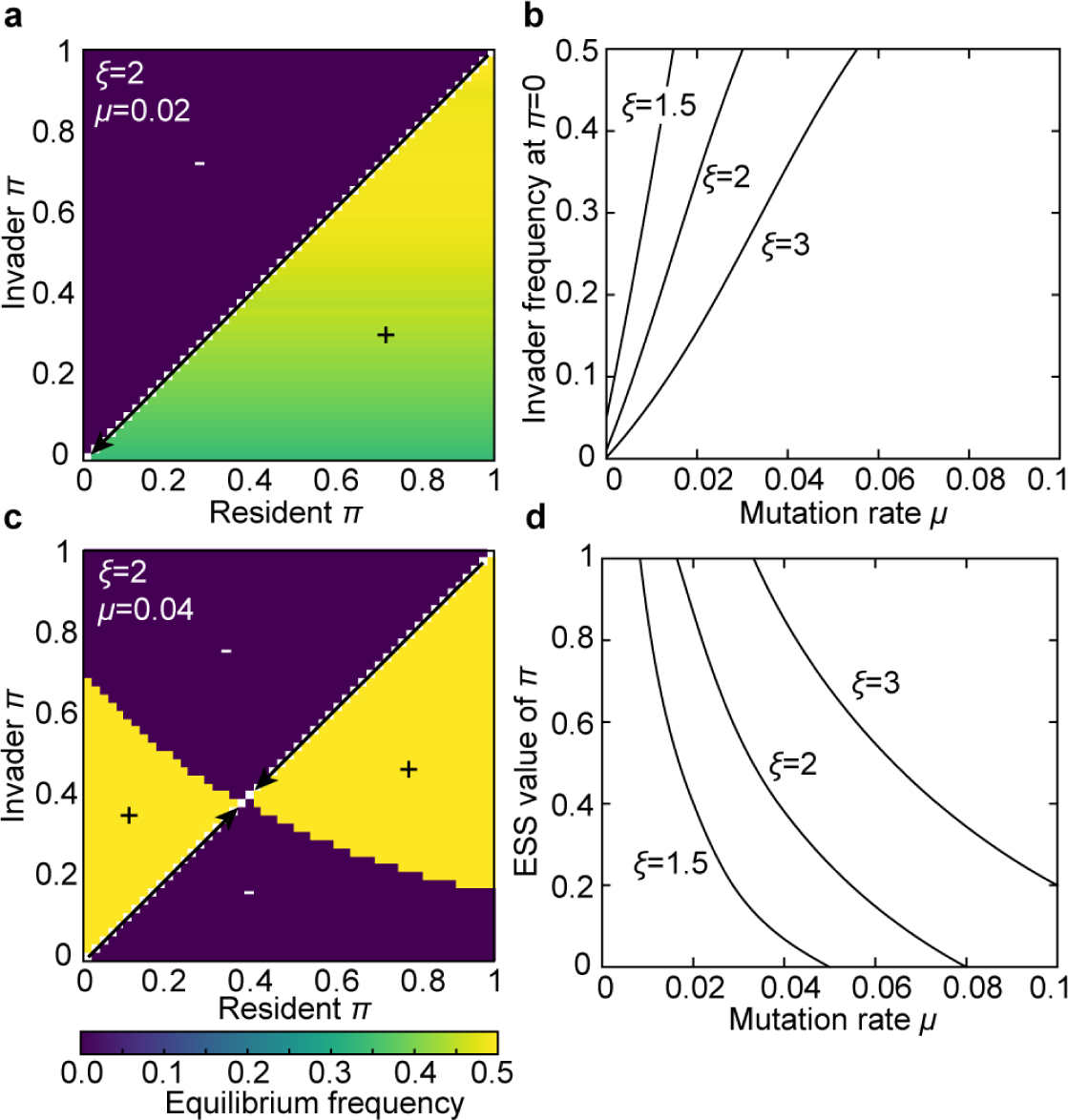
Evolutionary stability of asymmetric cytoplasmic inheritance in a population without mating types. Pairwise invisibility plot shows that with maternal control of cytoplasmic inheritance, the only asymmetric ESS is a strict UPI, i.e. *π*=0 (**a**), with the frequency of *U*_*m*_ at the ESS increasing with higher mutational load (**b**). In contrast, with paternal control the ESS can lie anywhere between the fully symmetric and strict uniparental inheritance (**c**), depending on mutation rates and epistatic interactions (**d**).

Under the paternal control of mitochondrial inheritance, the globally-attracting ESS lies between 0 ≤ *π*_ESS_ ≤ 1 (Fig. 3c), with the frequency of *U*_*p*_ allele always at *e*_2_ = 0.5. The deviation of *π* to either side of its *π*_ESS_ is detrimental: with *π* < *π*_ESS_ the asymmetry of mitochondrial inheritance increases, but the strength of long-term mitochondrial-nuclear associations is diminished, benefiting the resident population; similarly, *π* > *π*_ESS_ increases the strength of genetic linkage, but reduces the long-term effects of partial UPI—again to the detriment of the invader. The analysis further shows that the value of *π*_ESS_ goes down with increasing mutation rate *μ* and decreasing strength of epistasis *ξ* (Fig. 3d). The model therefore shows that limited mitochondrial mixing can be maintained in spite of its long-term fitness costs, but only if the inheritance-restriction allele causes the selective destruction of organelles inherited from the same mating type (paternal gamete by definition), and if the epistasis is negative.

## Invasion of self-incompatible mating-type alleles establishes the population-wide UPI

The above analysis shows that in a population without mating types the following equilibria with asymmetric inheritance of mitochondria are possible, depending on mutation rates and the mode of UPI nuclear regulation:

1) *U*_*m*_ at *e*_1_ < 0.5 with strict UPI *(π* = 0);
2) *U*_*m*_ at *e*_2_ = 0.5 with strict UPI *(π* = 0);
3) *U*_*p*_ at *e*_2_ = 0.5 with paternal leakage (0 ≤ *π* ≤ 1).

The last stable equilibrium for the allele *U*_*p*_ at *e*_1_ > 0.5 (Fig. 2a-b) is not considered here, as it cannot be reached from low allele frequencies. These equilibrium frequencies do not depend on the gamete mortality rate *δ*, as all gametes are compatible in the absence of mating types. While the frequency of uniparental mutants *U*_*m*_ or *U*_*p*_ can reach 0.5, the overall rate of asymmetric unions cannot exceed 0.5 as long as matings between identical gametes (*U*_*m*_×*U*_*m*_, *U*_*p*_×*U*_*p*_, *B×B*) are allowed. The biparental gamete unions can be eliminated, if a mutation in the intracellular signaling system leads to the incompatibility of cells carrying the same allele at the mitochondrial inheritance locus.

I next consider the invasion of self-incompatibility alleles *Mt*_1_ and *Mt*_2_, linked to either *U*_*m*_ /*U*_*p*_ or *B*. Due to the inherent symmetry of the system, the invasion of *U*_*m*_*Mt*_1_ under maternal inheritance regulation is in fact equivalent to invasion *BMt*_2_ under paternal control; similarly, the invasion of *U*_*p*_*Mt*_1_ is equivalent to the invasion of *BMt*_2_ under maternal regulation of cytoplasmic inheritance (Fig. 2a-b). It is therefore sufficient to study the evolution under maternal control, as the case of paternal regulation can be obtained by simply reversing the order of allele invasion, given that the initial equilibrium state exists and is stable under both modes of inheritance restriction.

With maternal allele *U*_*m*_ at equilibrium, the self-incompatible gamete class *U*_*m*_*Mt*_1_ invades replacing *U*_*m*_, but generally to a lower frequency than the ancestral *U*_*m*_ (Fig. 4a-b). Linkage to the mating type allele *Mt*_1_ ensures that symmetric unions between gametes carrying *U*_*m*_ become forbidden, increasing the frequency of uniparental matings and therefore leading to the higher long-term fitness advantage of *U*_*m*_ associated with more efficient removal of deleterious mutations. In a population consisting solely of *U*_*m*_*Mt*_1_ and *B*, all gamete unions involving *U*_*m*_*Mt*_1_ are therefore uniparental. At the same time, however, the subpopulation of *B* enjoys higher short-term fitness benefits of mitochondrial mixing in symmetric *B×B* unions, limiting the spread of the first mating type allele.

**Figure 4.**
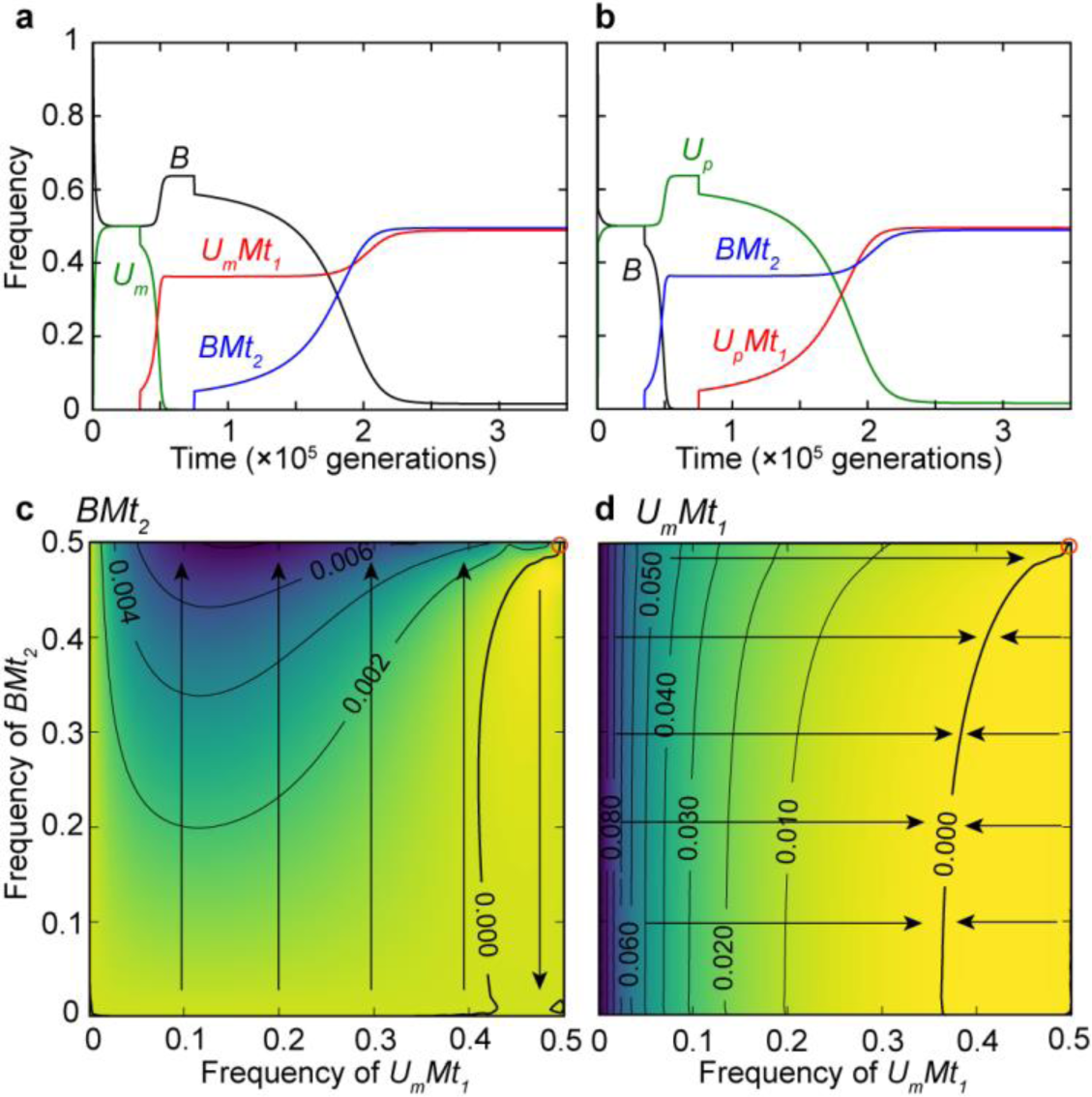
Invasion of self-incompatibility alleles *Mt*_1_ and *Mt*_2_ linked to the mitochondrial inheritance locus *U*_*m*_/*B* (*U*_*p*_/*B*). Due to the higher rate of uniparental gamete unions, *U*_*m*_*Mt*_1_ (or *BMt*_2_ assuming the paternal control) replaces the universally compatible allele *U*_*m*_ (*B*), but to a lower frequency (**a**-**b**). Similarly, the self-incompatible *BMt*_2_ (*U*_p_*Mt*_1_ under paternal regulation) then increases in frequency at the expense of *B* (*U*_*p*_), as it makes the asymmetric unions more frequent. The fitness advantage of the invading *BMt*_2_ (*U*_*p*_*Mt*_1_) only increases with its frequency (**c**). The fitness advantage of *U*_*m*_*Mt*_1_ (*BMt*_2_) over *B* (*U*_*p*_) also increases as *BMt*_2_ (*U*_*p*_*Mt*_1_) invades, as less frequent biparental gamete unions reduce the short-term fitness advantage of the universal resident *B* (**d**). Arrows indicate the expected direction of evolution in the population consisting of *U*_*m*_*Mt*_1_, *BMt*_2_ and *B*. Parameter values are *ξ* = 1.5, *μ* = 0.04, *π* = 0.4, *δ =* 10^−5^.

Similarly, in the invasion of the second self-incompatibility allele *Mt*_2_ linked to *B*, the gamete class *BMt*_2_ spreads at the expense of *B*, as it reduces the frequency of fully symmetric gamete unions and ensures that *BMt*_2_ inherits fit mitochondria from *U*_*m*_*Mt*_1_ uniparentally more often than the wild-type *B*. With the constant frequency of *U*_*m*_*Mt*_1_, the fitness advantage of *BMt*_2_ over *B* increases with its frequency (Fig. 4c). The spread of *BMt*_2_ also increases the long-term fitness advantage of the first invader *U*_*m*_*Mt*_1_ (Fig. 4d), as it reduces the short-term advantage of *B.* The spread of a second mating-type allele therefore reinforces the fitness advantage of the first, allowing both to reach high frequencies and even fix at low gamete mortality rates *δ.*

## Conditions favoring fixation of mating types

The two mating type alleles fix only if they are capable of replacing the universally compatible *U*_*m*_/*U*_*p*_ and *B*, which is opposed by two forces: 1) the short-term fitness advantage of mitochondrial mixing in subpopulations without mating types, and 2) the mating rate disadvantage of gametes that can mate only with a subset of the population. Indeed, the spread and fixation of *Mt*_1_ and *Mt*_2_ is favored by the factors increasing the long-term fitness effects of asymmetric transmission of mitochondria, and opposed by high gamete death rates *δ*.

First, regardless of the UPI regulation mode, the self-incompatible mating types replace the unisexual residents under weak epistatic interactions between mitochondrial mutations (*ξ* → 1, Fig. 5). The short-term fitness advantage of mitochondrial mixing in biparental gamete unions only applies to the case of negative epistasis between deleterious mutations (Radzvilavicius, 2016); at *ξ* = 1, for example, the uniparental invaders become universally advantageous, with their equilibrium frequency being limited only by the number of compatible mating partners. As the strength of negative epistatic interactions increases (*ξ* > 1), the short-term fitness effects of mitochondrial mixing become increasingly more important, opposing the spread of the two mating-type alleles. This effect can be partially alleviated in the presence of paternal leakage, as it tends to reduce the short-term fitness advantage of the universally-compatible gametes, favoring the spread of the two mating types (Fig. 5). The results therefore indicate the importance of paternal regulation of mitochondrial inheritance, as intermediate paternal leakage can be evolutionarily stable only under paternal control of cytoplasmic inheritance. Under strong epistasis and high levels of paternal leakage *π*, however, the first mating-type *U*_*m*_*Mt*_1_ fails to replace the universal *U*_*m*_, which subsequently prevents the invasion of *Mt*_2_ (Fig. 5).

**Figure 5.**
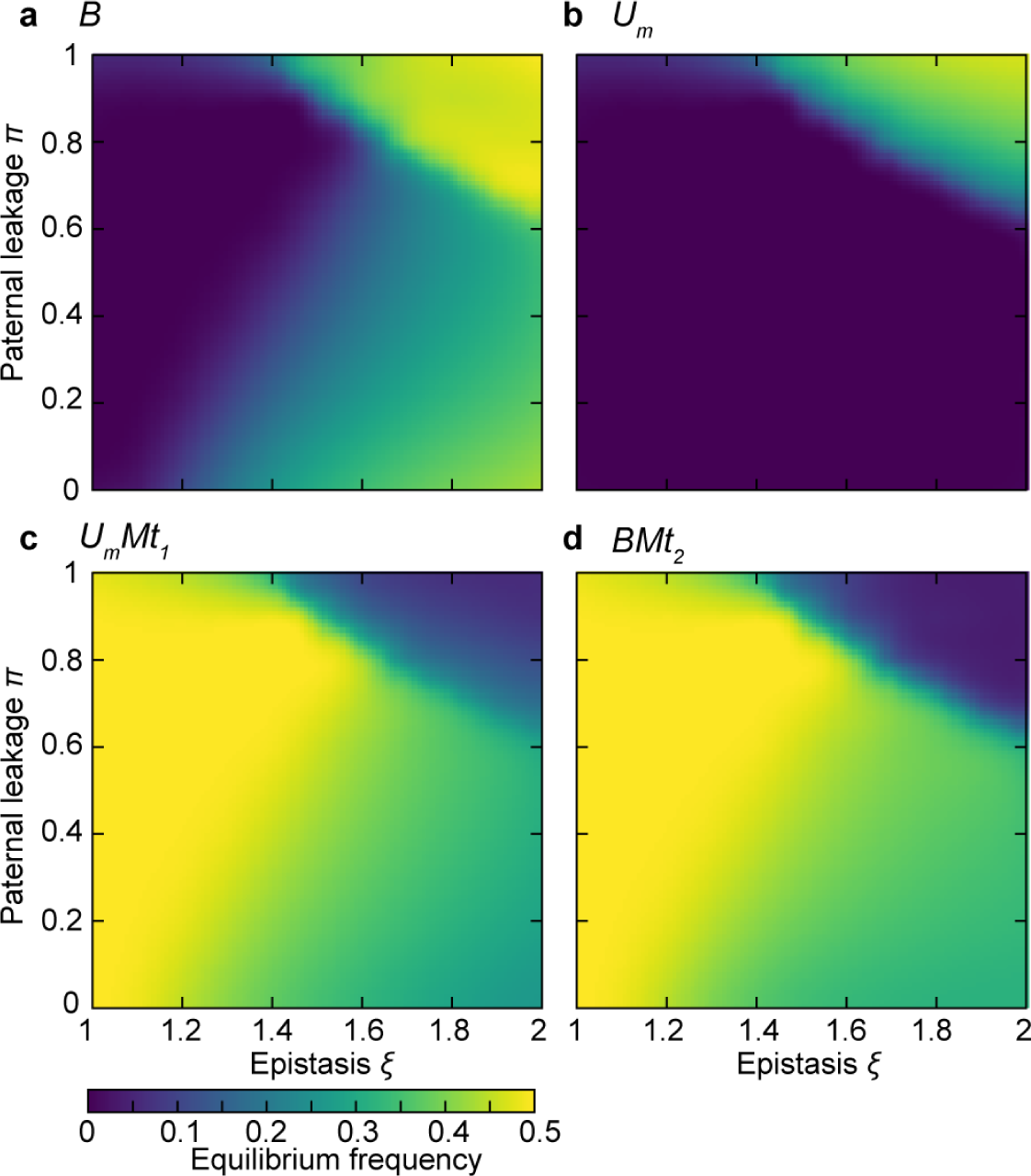
Equilibrium genotype frequencies for *B*, *U*_*m*_, *U*_*m*_*Mt*_1_ and *BMt*_2_ (*U*_*p*_, *B, BMt*_2_ and *U*_*p*_*Mt*_1_ under paternal inheritance regulation) after the invasion of both mating-type alleles linked to the mitochondrial inheritance loci, as a function of paternal leakage *π* and the strength of epistasis between mitochondrial mutations *ξ*. Self-incompatible mating types reach fixation under weak epistatic interactions, where the long-term fitness advantage of UPI is maximized. With stronger epistasis, the fixation of mating types is facilitated by the intermediate paternal leakage, implying the importance of paternal control of mitochondrial inheritance. Under strong negative epistasis and significant paternal leakage, mating types do not replace the universally compatible gametes due to weak long-term variance effects. Gamete death rate is set to *δ* = 0.001, the mitochondrial mutation rate is *μ =* 0.03.

High mitochondrial mutation rates favor the spread of mating-type alleles linked to the mitochondrial-inheritance locus (Fig. 6). This is consistent with the long-term effect of asymmetric mitochondrial transmission enhancing the efficacy of purifying selection against mitochondrial mutations due to higher variance in mitochondrial mutation load (Fig. 2). As expected, the evolutionary success of self-incompatibility alleles depends on the cost associated with the lower frequency of potential mating partners within the population, which is represented by the gamete mortality rate *δ* (Fig. 6). Under high mortality rates, gametes unable to find suitable mates are rapidly eliminated from the gamete pool; as a consequence, the universal gametes carrying *B* or *U*_*m*_/*U*_*p*_ prevail. With low gamete death rates *(δ* → 0), on the other hand, the self-incompatible mating types can persist longer, waiting for an encounter with a compatible mating partner, which favors their invasion and eventual fixation (Fig. 6). Indeed, while gamete self-incompatibility reduces the overall amount of potential mating partners within the population, with the evolution of mating types and mating-type associated pheromone-based intracellular attraction systems, gamete encounters are rarely random (Hadjivasiliou, 2015), and the part of the life cycle available for mate-finding could indeed be much longer than the time separating random encounters between mature reproductive cells.

**Figure 6.**
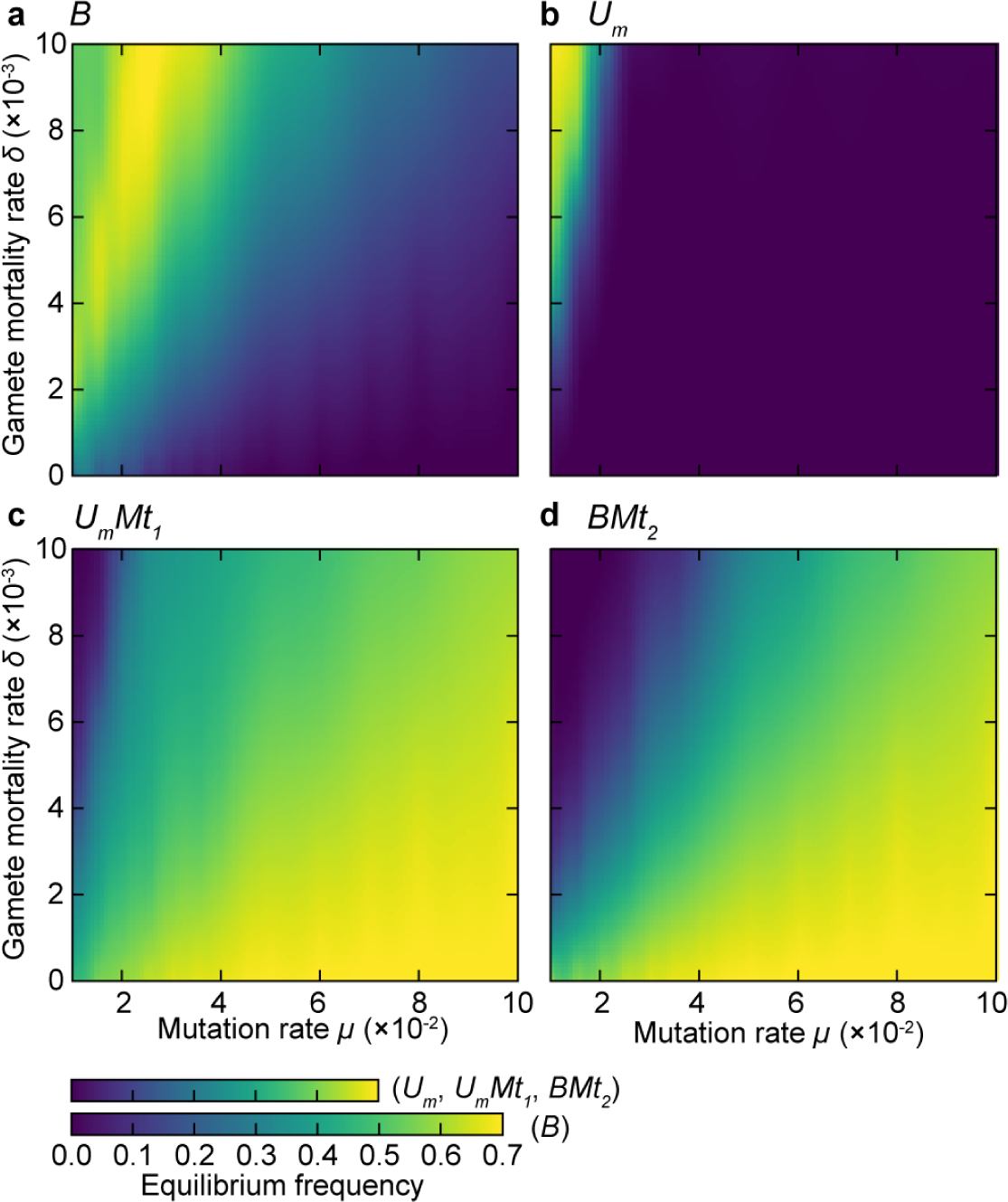
Equilibrium frequencies of genotypes *B, U*_*m*_, *U*_*m*_*Mt*_1_ and *BMt*_2_ (*U*_*p*_, *B, BMt*_2_ and *U*_*p*_*Mt*_1_ under paternal inheritance regulation) after the invasion of both mating-type alleles linked to the mitochondrial inheritance loci, as a function of gamete mortality rate *δ* and mitochondrial mutation rate *μ*. Self-incompatible gamete classes replace universal gamete types under high mutation rates and low gamete mortality rates. The strength of epistasis is *ξ* = 1.4, the rate of paternal leakage of mitochondria is set to *π* = 0.2.

## Discussion

The existence of two sexes or mating types is often regarded as an evolutionary conundrum, as both males and females are able to mate only with one-half of the population. The fitness benefits of gamete differentiation must therefore exceed the costs associated with the reduced mating opportunities. With the evolution of two mating types and the associated asymmetric signaling systems (Hoekstra, 1982; Hadjivasiliou et al., 2015), the encounters between gametes are not random, and the cost of being able to mate with only a part of the population might not be as high as assumed in the previous studies (Hadjivasiliou, 2013). I showed, that two mating-type alleles linked to the mitochondrial-inheritance locus can spread to fixation driven by the long-term advantage of asymmetric transmission of mitochondrial genes, which increases the efficacy of purifying selection against detrimental mitochondrial mutations, even though the invasion is opposed by the short-term fitness benefits of mitochondrial mixing in the biparental part of the population, and the limited availability of potential mating partners. Two mating types should therefore be expected to arise with fast accumulation of mitochondrial mutations, and high gamete-survival rates ensuring that the fitness cost associated with the need to locate a suitable mating partner is low. Additionally, the results show that paternal leakage of mitochondria can favor the evolution of mating types under negative epistasis, suggesting that paternal control of cytoplasmic inheritance could have been important.

Can the model account for the evolution of more than two gamete classes? Multiple mating types with uniparental inheritance of mitochondria are indeed known. For example, in true slime mold *Physarum polycephalum* with several mating types, uniparental inheritance of mitochondria via selective organelle digestion follows a complex hierarchy according to the allele at one of the mating-type loci (Moriyama and Kawano, 2003). Consider the invasion of a third self-incompatible mating type *Mt*_3_, linked to the mitochondrial inheritance locus with one of the existing alleles, say *U*_*m*_. The invader can therefore mate with both initial gamete classes, but part of the gamete unions will remain biparental (*U*_*m*_*Mt*_3_×*U*_*m*_*Mt*_1_). The model shows that *U*_*m*_*Mt*_3_ can invade only if it has a large initial mating-rate advantage, i.e. under gamete mortality rates ? high enough to compensate for the long-term fitness disadvantage due to less frequent uniparental unions. Interestingly, unions between certain pairs of gamete types in *Physarum polycephalum* are indeed biparental, indicating that some of the mating-type alleles might have arisen simply because of their mating-rate advantage (Moriyama and Kawano, 2003). Similar selective pressure has been suggested as a driving force for the evolution of multiple mating types in fungi without mobile gametes (Hurst, 1995).

The evolutionary stability of two mating types can therefore be explained by the high long-term fitness cost of biparental inheritance in the newly invading gamete class compared to the advantage of having more compatible mating partners. On the other hand, the results show that a third mating type allele invades much more readily in tandem with a new allele at the mitochondrial-inheritance locus *U*_*m1*_ coding for a novel mitochondrial recognition and destruction machinery. In this case, unions between all three gamete types remain uniparental throughout the invasion. Stable population of three mating types at equal frequencies can therefore become established even at low gamete mortality rates *δ*; the same remains true for subsequently invading mating-type alleles. Under these assumptions the number of mating types in the population would seem to be limited only by the amount of distinct molecular mechanisms controlling the transmission of organelle genomes. This scenario, however, is critically dependent on the simultaneous invasion of two novel mutations at linked loci, both controlling vastly complex processes of mate recognition and mitochondrial destruction, and is therefore highly unlikely.

How does the present analysis compare to the previous work suggesting that mating types could have evolved together with the uniparental inheritance of cytoplasmic genes? In the model of Hurst and Hamilton (1992), gamete classes evolve to eliminate the conflict between cytoplasmic genes inherited from distinct parents (Hurst, 1995). Similarly, Hutson and Law (1993) considered the spread of “selfish” endosymbionts with fixed fitness costs. While multiple cases of selfish organelles and genetic elements are known (Schable and Wise, 1998; Taylor et al., 2002; Clark et al., 2012), it is unlikely that they occur at high enough frequencies to account for the universality of mating types with uniparental transmission of cytoplasm in eukaryotes. Selection against deleterious mitochondrial mutations, on the other hand, provides a more general explanation. While the rates of mtDNA mutation accumulation vary substantially between groups, in many cases they can exceed the evolution rates of nuclear genes by an order of magnitude, supporting the hypothesis (Lynch, 2006). Given the central role of mitochondria in eukaryotic metabolism, it is not surprising that mechanisms facilitating removal of deleterious mitochondrial mutations are selected for, e.g. germline bottlenecks or early germline sequestration in bilaterians. The evolution of two mating types with uniparental organelle transmission might therefore be a consequence of the fundamental requirement for high mitochondrial quality in complex eukaryotes.

